# Impact of variants of concern on SARS-CoV-2 viral dynamics in non-human primates

**DOI:** 10.1101/2022.11.09.515748

**Authors:** Aurélien Marc, Romain Marlin, Flora Donati, Mélanie Prague, Marion Kerioui, Cécile Hérate, Marie Alexandre, Nathalie Dereuddre-bosquet, Julie Bertrand, Vanessa Contreras, Sylvie Behillil, Pauline Maisonnasse, Sylvie Van Der Werf, Roger Le Grand, Jérémie Guedj

## Abstract

The impact of variants of concern (VoC) on SARS-CoV-2 viral dynamics remains poorly understood and essentially relies on observational studies subject to various sorts of biases. In contrast, experimental models of infection constitute a powerful model to perform controlled comparisons of the viral dynamics observed with VoC and better quantify how VoC escape from the immune response.

Here we used molecular and infectious viral load of 78 cynomolgus macaques to characterize in detail the effects of VoC on viral dynamics. We first developed a mathematical model that recapitulate the observed dynamics, and we found that the best model describing the data assumed a rapid antigen-dependent stimulation of the immune response leading to a rapid reduction of viral infectivity. When compared with the historical variant, all VoC except beta were associated with an escape from this immune response, and this effect was particularly sensitive for delta and omicron variant (p<10^−6^ for both). Interestingly, delta variant was associated with a 1.8-fold increased viral production rate (p=0.046), while conversely omicron variant was associated with a 14-fold reduction in viral production rate (p<10^−6^). During a natural infection, our models predict that delta variant is associated with a higher peak viral RNA than omicron variant (7.6 log_10_ copies/mL 95% CI 6.8 – 8 for delta; 5.6 log_10_ copies/mL 95% CI 4.8 – 6.3 for omicron) while having similar peak infectious titers (3.7 log_10_ PFU/mL 95% CI 2.4 – 4.6 for delta; 2.8 log_10_ PFU/mL 95% CI 1.9 – 3.8 for omicron). These results provide a detailed picture of the effects of VoC on total and infectious viral load and may help understand some differences observed in the patterns of viral transmission of these viruses.

## Introduction

The sever acute respiratory coronavirus 2 (SARS-CoV-2) is the causative agent of the Coronavirus-induced disease 2019 (COVID-19) cumulating more than 500 million cases and over 18 million death as measured by excess mortality as the end of 2022 (1,2). Repeatedly, several variants have emerged and although most of them vanished quickly, some of them, called Variants of Concern (VoC), in particular alpha, beta, gamma, delta and omicron have caused dramatic epidemic rebounds (3–5). These variants have acquired specific mutations enhancing their infectious capacities and escaping the immune response, leading to a dramatic loss of efficacy of monoclonal antibodies (6). They have also caused a large drop in vaccine efficacy against disease acquisition even though until now vaccine remain largely effective against severe disease (7–9).

While several millions of individuals have been infected by these VoC, we still do not have a precise understanding on the effects of VoC on viral load. Even though some effects on larger levels of viral excretion have been reported (10–13), these studies often lack of robustness, and may be biased by many confounding factors that complicate comparisons, in particular reporting biases, heterogeneity in the incubation period and vaccination coverage.

In that context where human clinical data are difficult to interpret, the non-human primate (NHP) experimental model offers a unique opportunity to describe infection with SARS-CoV-2 in detail in a fully controlled environment. Since 2020, our group has conducted many studies to evaluate the effects of antiviral drugs or vaccines in this model (14,15), and showed its large predictive value (16). Here, we analysed retrospectively viral load data obtained in 78 animals that were included as control arms of these studies and that were infected with different strains of SARS-CoV-2 (historical, beta, gamma, delta and omicron (BA.1)). In addition, we performed longitudinal measures of viral culture to evaluate a potential effect of VoC on viral infectivity. Using the techniques of mathematical modelling, we characterize the viral kinetics in these animals and we discuss their biological insights.

## Results

### Variant of concern viral kinetics

Several biomarkers were measured, both genomic RNA and subgenomic RNA were quantified at regular interval over all the study period and infectious titers at 2 times points. All macaques developed a rapid infection with genomic viral load peaking between 2- and 3-day post-infection (dpi) for the historical and beta variant, 3.5 dpi for variant delta and 4 dpi for variants gamma and omicron (BA.1). Genomic viral load was cleared at 8 dpi for the historical variant, 10 dpi for the beta variant, at 12 dpi for variants delta and omicron (BA.1) and at 14 dpi for variant gamma (Fig 1 and S1 Table). In addition to viral RNA, infectious titers were measured for 41 animals. Infectious titers were measured by Tissue Culture Infectious Dose (TCID_50_) from nasopharyngeal swab sampled at 2 time points per animal (day 2, 3 or 4 plus at day 5 or 7 post-infection). As we included several control animals from different studies, infected with either TCID_50_ or Plaque Forming Units (PFU), all TCID_50_ were converted to PFU assuming 1 PFU = 0.7 TCID_50_ (17). All infectious titers quickly dropped to undetectable levels for the historical variant at 5 dpi, where for the other variants the infectious titers remained consistent over the course of the infection (Fig 1).

**Fig 1.**
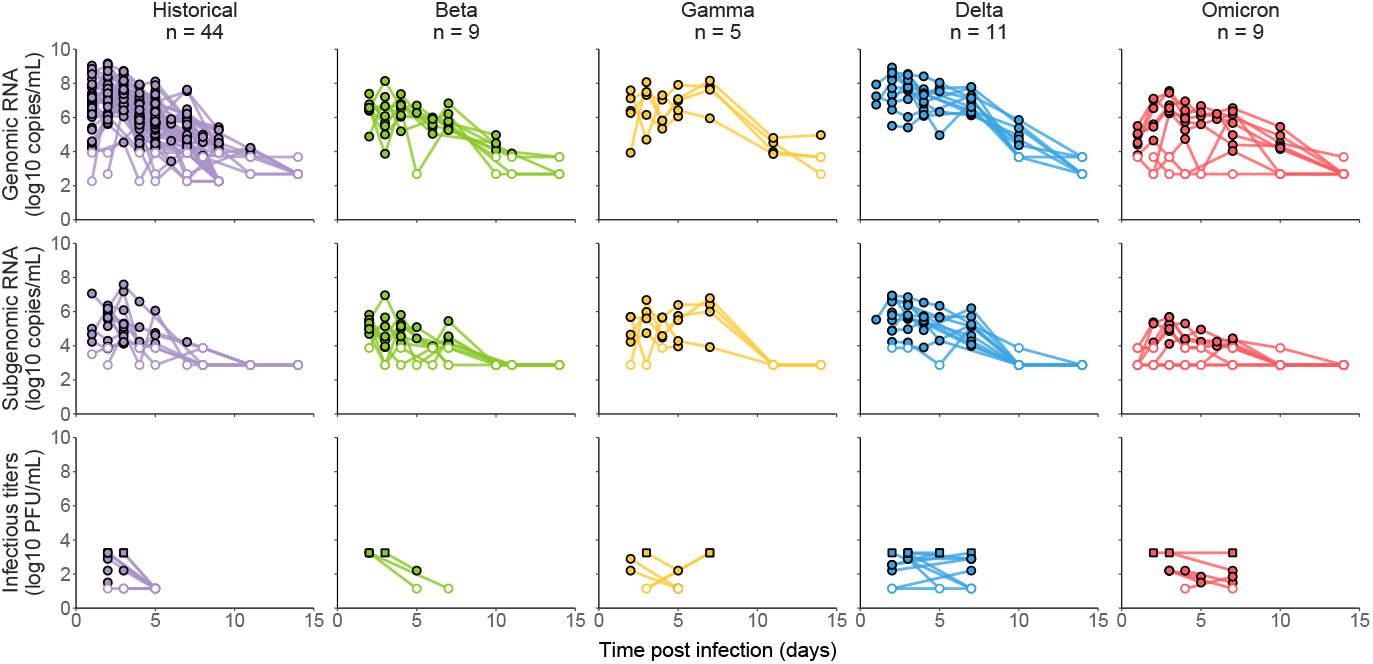
Longitudinal measurements of genomic RNA, subgenomic RNA and infectious titers in 78 infected cynomolgus macaques. Both limit of quantification and detection are depicted as empty dots, the latter being lower. Upper limit of detection is depicted as filled squares.

### Viral dynamic model

To account for the quick drop in infectious titers observed in the historical variant, (Fig 1 and S1 Fig) several models incorporating an action of an antigen-mediated immune response were tested (Fig 2). All models, except a model targeting the viral production parameter, provided an improvement of BIC compared to a target cell limited model (Table 1). We found that a model targeting the infectious ratio best described our data. In the following, we discuss the parameter values of the final constructed model accounting for both an effect of the immune effector and variant specific effect on the parameters (see below). For the historical variant, we estimated the infectivity rate parameter *β*at 1.86×10^−5^ copies^-1^.d^-1^ (95% confidence interval (CI) 1×10^−5^ – 3.39×10^−5^) and the loss rate of infected cells *δ* at 1.38 d^-1^ (95% CI 1.22 – 1.55), corresponding to a half-life of 12 hours. We estimated the viral load production parameter *p* at 9.44×10^5^ copies.cells^-1^.day^-1^ (95% CI 2.1×10^5^ – 1.68×10^6^). This corresponds to a within-host basic reproductive number *R*_0_ (i.e., the number of newly infected cells by one infected cell at the beginning of the infection) of 3.1 (95% CI 2 – 4.3) and a burst size (i.e the total number of infectious virus produced by one cell over its lifespan at the beginning of the infection) of 136 (95% CI 121 - 153).

**Table 1:** Alternative immune response models.

| Models | Description | ΔBIC |
| --- | --- | --- |
| Reference model | Absence of immune response | — |
| **Model 1** | **Reduction of the infectious ratio** | **−42.8** |
| Model 2 | Increase in infected cell clearance | −14.8 |
| Model 3 | Reduction of viral infection rate | −36.1 |
| Model 4 | Reduction of the viral production | +9 |

**Fig 2.**
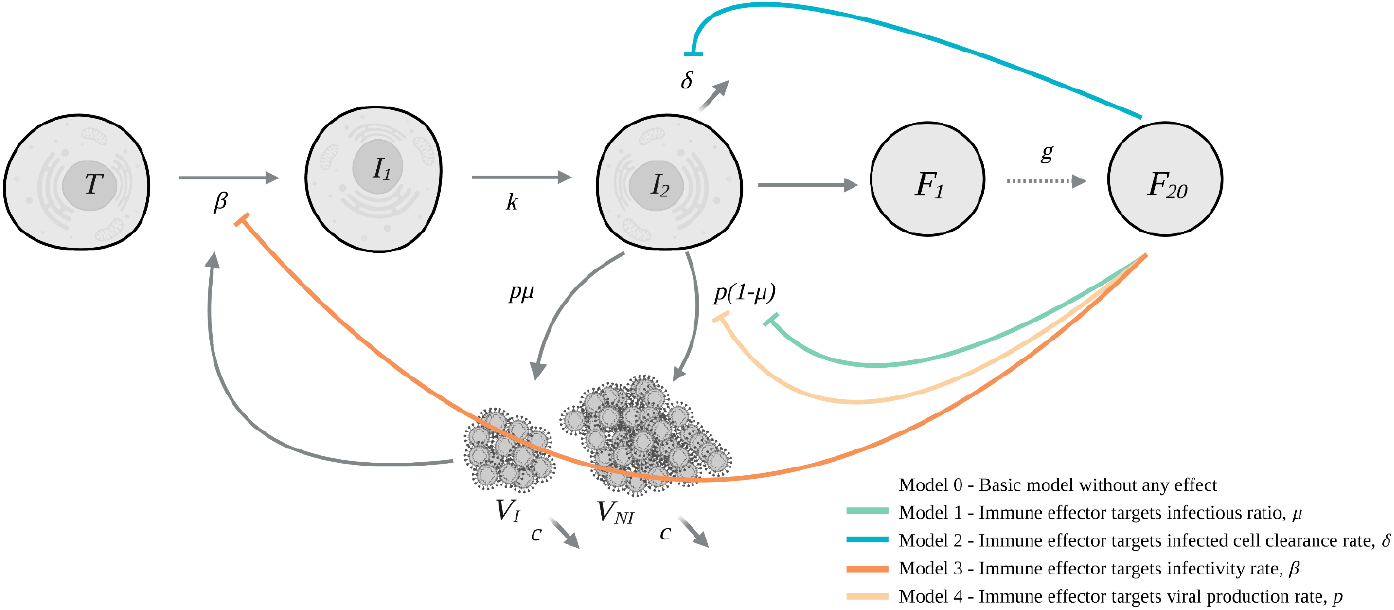
Schematic model of SARS-CoV-2 infection and action of the immune system. The basic model is a target cell limited model without any immune response. The parameters are : ***β*** the infectivity rate, ***k*** the transfer rate between non-productive and productive infected cells, ***δ*** the loss rate of productive infected cells, ***p*** the viral production rate, ***µ*** the ratio of infectious virus, ***g*** the transfer rate between the compartments of the immune response and ***c*** the loss rate of both infectious and non-infectious virus

### VoC specific effect on viral dynamic parameters

Once an effect of the immune response was selected, a covariate search algorithm was used to find the most likely VoC associated effects (see methods) and considered the historical variant as the reference. Several variant-specific covariates were found on viral kinetics parameters that we detail below (Fig 3 and S2 Table). First, beta variant was characterized with a reduced infected cells death rate (*δ*) by a factor of 0.7 (95% CI 0.6 − 0.9) compared with the historical variant (p-value < 0.01). This led to an infected cell half-life of 17 hours and resulted in a longer period of viral load shedding as infected cells produced viruses for longer period of time. Gamma variant had an effect on the parameter *θ* (p-value < 0.001), the amount of immune effector *F*_20_ required to reduce by half the infectious ratio, increasing it by a factor of 9508 (95% CI 387 − 50 041) resulting in higher peak viral load and a longer duration of infectious virus shedding (Fig 4). Variant delta is characterized by an effect on both *θ* (p-value < 0.001) and the viral production parameter *p* (p-value < 0.05), increasing those parameters by factors 336 (95% CI 49 − 1191) and 1.78 (95% CI 1 − 3) respectively. Finally, omicron variant (BA.1) affected the parameters of the immune system *θ* (p-value < 0.001), the viral production rate parameter *p* (p-value < 0.001) and the infectious ratio *µ* (p-value < 0.001) modifying them by factors 229 (95% CI 27 − 884), 0.07 (95% CI 0.02 − 0.2) and 18 (95% CI 4 − 51) respectively (Fig 4). The model well reproduced the viral load of all animals in the individuals fits (S2 Fig). Additionally, we performed a sensitivity analysis on our best model (i.e. Model 1 including an effect on the infectious ratio *μ*). We tested several delays of the immune effector (from 1 to 6 days post infection) and several numbers of transfer compartments (from 5 to 30) and performed the covariate search on all models. We found that a delay of 3 days yielded the best results (S3 Table) and very similar covariate were selected across all models (S3 and S4 Fig).

**Fig 3.**
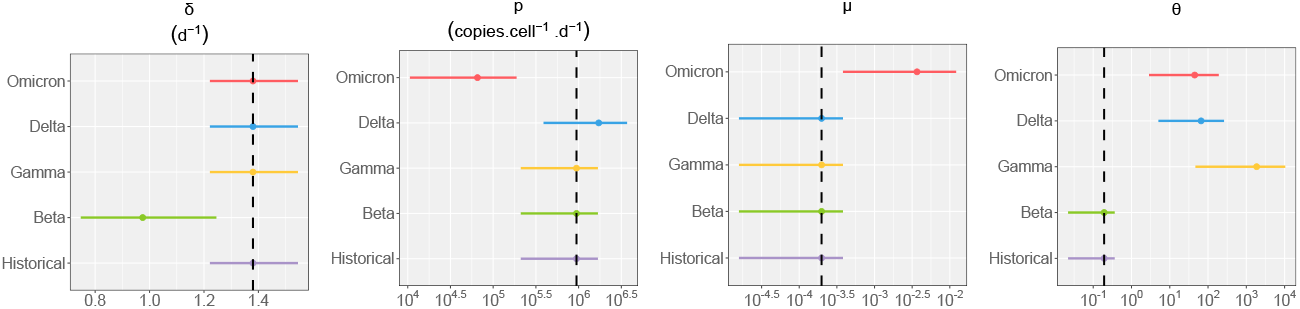
Estimated population parameters for each variant. We represent the mean value and 95% confidence interval of populations parameters for each variant. We represent only parameters having at least one variant-specific effect. Full table for population parameters is in S2 Table. The dashed black line represents the historical value.

**Fig 4.**
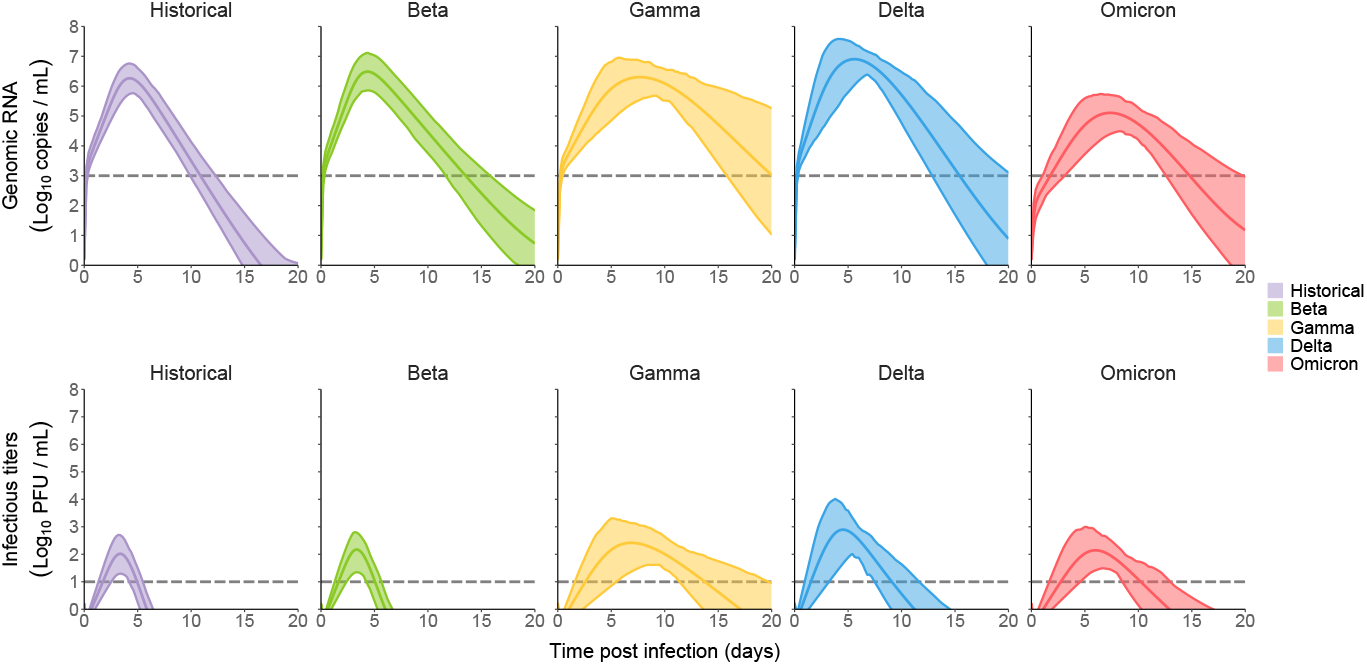
Simulation of variant of concern impact on viral load. Using simulations, we sampled parameters considering both the uncertainty in the estimation and the inter-individual variability (see methods). We represent the mean viral load of all variants and its 95% confidence interval. Dotted lines are the limits of detections

### Predicted impact of variants in a natural infection setting

The main limitation of translating these results to humans is the fact that infection in animals is done with a large inoculum dose (10^5^-10^6^ PFU), while human infections are presumably initiated with much lower virus dose (18). Human experimental infections were performed with 10 TCID_50_ (19) in the nose, i.e., 10,000-100,000 times less virus than in the animal model. Using simulations with lower inoculum, considering both uncertainty in the estimation and inter-individual variability (see methods), we are able to derive metrics of interest for each variant.

The historical variant is characterized by a mean time to peak of 4.3 dpi (95% CI 3.7 − 4.8) and of 3.5 dpi (95% CI 3 − 3.9) for genomic RNA and infectious titers respectively. We found a mean peak viral load of 6.3 log_10_ copies/mL (95% CI 5.5 − 7) and of 2.1 PFU/mL (95% CI 1.2 − 2.9) for genomic RNA and infectious titers, respectively.

The reduced infected cell clearance rate of the beta variant resulted in a longer period of viral load shedding. The duration of the acute infection stage was consequently increased from 10.9 days (95% CI 9.5 – 13.1) for the historical variant to 13.4 days (95% CI 11.1 – 15.7) for the beta variant.

All variants except beta have shown an effect on the antigen-mediated response, greatly reducing its impact on viral kinetics. As the effect of the antigen-mediated response was reduced, the infectious ratio was increased leading to more infectious particles produced over longer periods of time. This led to the increase of the infectious titers clearance stage duration from 1.5 days for the historical variant (95% CI 0.6 – 1.9) to 6 days (95% CI 4.4 – 7.5), 3.8 days (95% CI 3.1 - 4.6) and 3.7 days (95% CI 2.8 – 4.5) for the gamma, delta and omicron variants respectively (Fig 5). This is in line with numbers of studies showing the immune escape capabilities of those variants (20–22).

**Fig 5.**
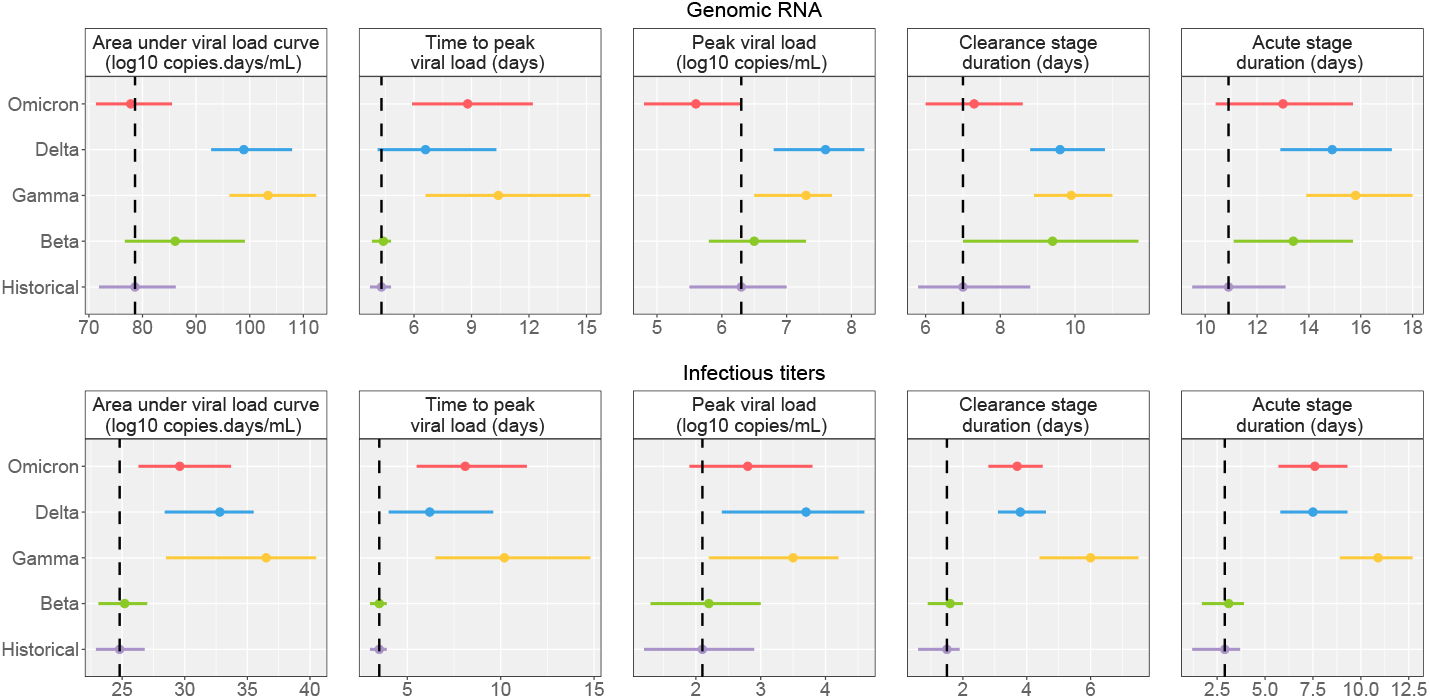
Impact of VoC on viral load metrics in the context of an infection with a low inoculum. We represent the mean and 95% confidence interval for each variant. The dashed black line represents the historical mean value.

An effect increasing the viral production parameter (*p*), as observed for the delta variant, results in largely higher peak viral load of 7.6 log_10_ copies/mL (95% CI 6.8 – 8.2) and peak infectious titers of 3.7 PFU/mL (95% CI 2.4 – 4.6). Conversely, an effect reducing the viral production parameter, as observed for the omicron variant, results in lower peak viral load compared to the historical variant of 5.6 log_10_ copies/mL (4.8 – 6.3) but very similar peak infectious titers at 2.8 PFU/mL (95% CI 1.9 – 3.8). This is due to an effect of omicron on the infectious ratio, increasing the proportion of infectious virus produced.

## Discussion

Here, we used mechanistic models to characterize in detail the viral dynamics of the main variants of concern in an experimental model of non-human primates. We evaluated the impact of an antigen-mediated immune response on the viral dynamics and found that an effect reducing the infectious ratio best described our data. Some of the variants of concern, gamma, delta and omicron (BA.1) showed a strong ability to escape this response greatly increasing the number of infectious viruses produced over the course of the infection compared to the historical variant. Interestingly, the delta variant was associated with an increased viral production rate, whereas the omicron variant was associated with a lower viral production rate but a higher infectious ratio.

Using simulations in a natural infection scenario, we found that omicron infections, relative to delta infections, are associated with lower peak viral RNA and reduced duration of viral RNA clearance while having similar peak infectious titers and duration of infectious titers clearance.

These results suggest that omicron’s infectiousness cannot be attributed to an increased viral RNA production but maybe due to an immune escape coupled with an increased infectious ratio, greatly increasing the number of infectious particles produced.

Although many other factors are at play to explain the increased transmissibility of certain variants of concern, differences in viral dynamics can provides insights into the biology of those variants. As such, delta infections featuring increased peak viral load and infectious titers can increase the risk of “superspreading” events and infections outside of close-contact settings (23). Omicron infections, on the other hand, featuring lower peak viral load concentration (24) but similar infectious titers respective to other variants, may result in transmission events that would not occur with other variants because insufficient infectious titers would be produced. These results are coherent with reports showing lower pathogenicity of omicron infection (25), as they are associated with lower viral burden.

The combination of immune escape abilities, increased infectious ratio and longer duration of infectious virus shedding could be a possible mechanism to explain the enhanced transmissibility of omicron variant. As such, the quantification of infectious titers over time is crucial to inform further public health policies and adjust the isolation period accordingly.

Our study has some important limitations. First, although we can characterise in detail the viral dynamics of SARS-CoV-2 in nonhuman primates in a controlled environment, the inoculated dose is extremely high (10 000 to 100 000 times higher (19)) compared with human infection. This leads to rapid saturation of target cells and makes it difficult to accurately estimate the early phase of infection. In the future, studies evaluating lower inoculum in NHP can greatly improve the precision in the estimation of the early phase of infection. Second, we developed an extension of the target cell limited model considering the effect of an antigen-mediated immune response decreasing the infectious ratio *µ*. We here attribute this effect to the immune system but we have no information to which immune effectors (antibodies, cytokines, cytotoxic cells, natural killers, intracellular processes etc…) this could be linked if even attributable to one. This type of antigen-mediated response allows us to incorporate the effect of time on a parameter but the underlying biological mechanisms are unclear and may be due to inherent differences between variants not captured by any covariates.

Third, we assumed a 3-days delay in the establishment of this antigen-mediated reduction of the infectious ratio and verified that it performed best in a sensitivity analysis (S4 Fig). Although there is some variability, the covariates search is overall consistent.

Fourth, the infectious titers are only a measure of *in vitro* infectivity, and to what extent they translate into infectiousness is unknown. In addition, both the upper and lower limit of quantification makes it difficult to precisely estimate the infectious ratio parameter *µ*. Finally, in a context where more than half of the world population has received at least one dose of COVID-19 vaccine (26), there is very little information on the natural infection with different variants. Additional data with vaccinated animals could help differentiate certain aspects of the abilities of the new variants to escape the immune system.

## Materials and methods

### Experimental procedure

Data comes from studies performed on cynomolgus macaques to evaluate the viral dynamics of SARS-CoV-2 variants. Our study includes 78 cynomolgus macaques (*Macaca fascicularis*) coming from control arms of several studies and have received no pharmacological interventions besides placebo. All animals were infected with doses ranging from 7×10^4^ to 10^6^ PFU of different SARS-CoV-2 strains. Animals are infected via both nasopharyngeal and intratracheal route with 10% of the initial volume administered in the nose and 90% in the trachea. The study is composed of 5 groups, each infected with a different SARS-CoV-2 strains: 44 Historical (hCoV-19/France/lDF0372/2020 strain; GISAID EpiCoV platform under accession number EPI_ISL_406596), 9 Bêta (B.1.351 - hCoV-19/USA/MD-HP01542/2021, BEI NR-55283), 5 Gamma (P.1 - hCoV-19/Japan/TY7-503/2021, BEI NR-54984), 11 Delta (B.1.617.2 - hCoV-19/USA/MD-HP05647/2021, BEI NR-55674) and 9 Omicron (B.1.1.529 – hCoV-19/USA/MD-HP20874/2021, BEI NR-56462). For each group both genomic RNA and subgenomic RNA swab samples were quantified using real time PCR in both the nasopharynx and in the trachea. For 41 animals (13 Historical, 3 Beta, 5 Gamma, 11 Delta and 7 Omicron (BA.1)) infectious titers were measured at 2 time points, early (2, 3 or 4 days post infection) and late (5 or 7 days post infection) using Tissue Culture Infectious Dose (TCID_50_) from nasopharyngeal swab samples (16). As we included animals from different studies that were inoculated with different methods (PFU or TCID_50_), we normalized all measures of infectious titers by converting all TCID_50_ measurements to Plaque Forming Units (PFU) using the formula 1 PFU = 0.7 TCID_50_ (17). As no infectious titers were measured in the trachea samples, we focused the main analysis on the nasopharyngeal compartment. The results mainly focus on the genomic viral load as the subgenomic is a directly proportional to the latter.

### Basic viral dynamic model

We used a previously described model of SARS-COV-2 viral dynamics to reconstruct the nasopharyngeal viral load of infected animals. In this model, target cells (T) become infected cells (I_1_) at a rate *β*. Infected cells transition into productive infected cells (I_2_) at a rate *k* and produce infectious virus (V_I_) at a rate *pµ* and non-infectious virus (V_NI_) at a rate *p*(1 − *µ*). Productive infected cells are cleared at a rate *δ* and both infectious and non-infectious virus are cleared at a rate *c*. The basic within-host reproductive number, representing the number of newly infected cells by one infected cell, is 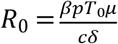 and the burst-size, representing the number of infectious virus produced by on infected cells over its lifespan, is 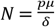. The model is described with the following set of ordinary differential equations:

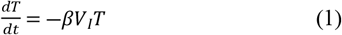

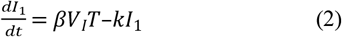

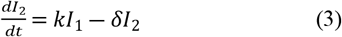

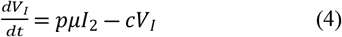

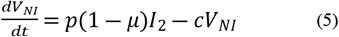

### Assumption on parameter values

Some parameters of the model were fixed to ensure identifiability. The transfer rate parameter between infected cells and productive infected cells was fixed to *k* = 4 day^-1^ (corresponding to a mean duration of the eclipse phase, i.e. the time for infected cells to start producing viruses, of 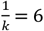 hours) (27). The viral clearance c was set to 10 day^-1^ based on previous work (14,16,28). As only the product *pT*_0_ is identifiable, we choose to fix the initial number of target cell to *T*_0_ = 12 500 cells following the same assumptions as in (16). As the nasal cavity of the animals is small, a substantial fraction of the inoculum does not penetrate the upper respiratory tract. To account for this, we introduced a parameter *h* representing the proportion of the inoculum that arrive on the site of infection. We fixed this parameter at 20% with a standard deviation of 20% to allow for individual variability. As both the initial infectious inoculum and the number of RNA copies were known we used that information as our initial condition for the infectious virus and non-infectious virus compartment. Therefore, our initial conditions were set to :

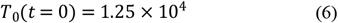

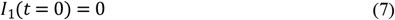

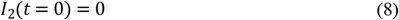

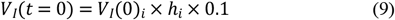

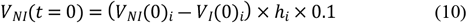

Where *V*_*I*_(0)_*i*_ is the administered dose in PFU of subject *i, V*_*NI*_(0)_*i*_ is the total number of RNA copies in the initial inoculum of subject *i* and *h*_*i*_ is the proportion of the inoculum actively initiating the infection.

### Models incorporating antigen-mediated immune response

To account for the quick drop in infectious titers observed for the historical variant (Fig 1 and S2 Fig), we tested several models incorporating an action of an antigen-mediated immune response. We assumed a delay of 3 days for the immune response to take place to account for the differentiation and proliferation of the immune response (29). We modelled this delayed immune effector compartment using the Linear Chain Trick (LCT) assuming an Erlang distribution with *j* = 20 transition compartment and a mean time spent in those compartment of *τ* = 3 d^-1^ (30). This number of compartments allowed us to shift the distribution of the time spent in the transition’s states from an exponential to a normal distribution. The equations for the transfer compartments are written as follows:

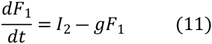

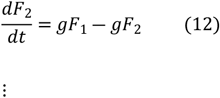

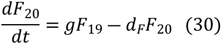

In the following only the compartment *F*_20_ will serve as the effector for the action of the immune system. The transfer rate parameter *g* is then written as 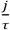 and fixed to 6.67 d^-1^ and the loss rate of the final effector *d*_*F*_ is fixed to 0.4 d^-1^ (28). Several modes of action of the response system were tested:

#### Model 1 : Immune effector decreases the infectious ratio *µ*

In this model, the immune effector directly decreases the infectious ratio parameter *µ* using an Emax function type expression :

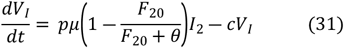

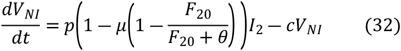

With *θ* being the amount of immune effector *F*_20_ needed to reduce by half the infectious ratio.

#### Model 2 : Immune effector increases infected productive cells death rate *δ*

The death rate of infected cells is increased in proportion to the amount of immune effector *F*_20_.

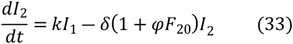

Where *ψ* is the strength of the immune system.

#### Model 3 : Immune effector reduces the infectivity rate *β*

In this model, the immune effector blocks virus entry in the cells by reducing the infectivity parameter *β*.

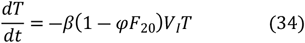

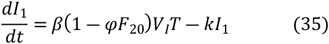

#### Model 4 : Immune effector reduces the production rate *p*

In the same way as model 1, the viral load production parameter is reduced by the immune effector with an Emax type function:

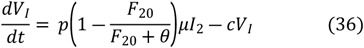

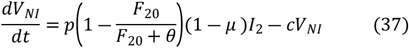

All models were compared based on the Bayesian Information Criterion (BIC). We selected the model that yielded the lowest BIC and the best individual fits.

### Statistical model

Parameter estimation was performed using non-linear mixed effect modelling. The statistical models describing the genomic RNA, subgenomic RNA and the infectious titers are:

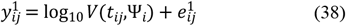

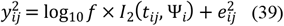

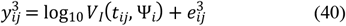

Where the superscript 1, 2 and 3 refers to the genomic RNA, subgenomic RNA and infectious titers, respectively. We denote *y*_*ij*_ is the *j*^*th*^ observation of subject *i* at time *t*_*ij*_, with *i* ∈ 1, …, N and *j* ∈ 1, …, *n*_*i*_ with N the number of subject and *n*_*i*_ the number of observations for subject *i*. The function describing the total viral load kinetics *V*(*t*_*ij*_, *Ψ*_*i*_) predicted by the model at time *t*_*ij*_ defined as: *V*_*I*_(*t*_*ij*_, *Ψ*_*i*_) + *V*_*NI*_(*t*_*ij*_, *Ψ*_*i*_) predicted by the model at time *t*_*ij*_. The The vector of individual parameters of subject *i* is noted *Ψ*_*i*_ and *e*_*ij*_ is the additive residual Gaussian error of constant standard deviation *σ*. The vector of individual parameters depends on a fixed effects vector and on an individual random effects vector, which follows a normal centered distribution with diagonal variance-covariance matrix *Ω*. All parameters follow a log-normal distribution to ensure positivity except both parameters *µ* and *h* which follows logit-normal distribution and are bounded between 0 and 1. We assumed random effect on all parameters and removed them using backward procedure, if they were < 0.1 or their RSE > 50%. All biomarkers (i.e. genomic RNA, subgenomic RNA and infectious titers) were fitted simultaneously.

### Selection of variant-specific effect on the viral dynamic parameters

Using the best model selected at the previous step, we sought to identify VoC-specific effect on the parameters of the model (*β, δ, p,µ* and *θ*). We first performed a backward selection of the random effects removing non-significant ones (i.e. relative standard error > 50%) if the BIC wasn’t degraded by more than 2 points. We then used the Conditional Sampling use for Stepwise Approach on Correlation tests (COSSAC) to identify variant specific effect (31). Then a backward procedure was used to remove any non-significant covariate effect with a Wald test (i.e. the covariate was removed if its coefficient effect relative standard error was > 50%). This procedure was repeated until all nonsignificant covariate effects had been eliminated. Additionally, we performed a sensitivity analysis on our best structural model. We tested for several delays in the establishment of the antigen-mediated effector (from 1 to 6 days) and on the number of transitions compartments (from 5 to 30) and then performed the covariate search on all model combinations.

### Simulation of natural human infection

Finally, we used our final model to assess the impact of variants of concern on viral load and viral infectivity in a natural infection setting. We used a starting inoculum of 10 infectious virus, as described in an experimental challenge conducted in England (19) to simulate a human infection. The initial conditions are then written as:

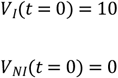

We provided confidence interval on the mean predicted viral load, considering both the uncertainty in the estimation and the inter-individual variability. We first sampled M = 100 population parameters in their estimation distribution and then, for each variant, sampled N = 30 individual parameters from each sets of population parameters (leading to 3000 individual parameters per variant). We calculated the predicted viral load of all individuals and derived the mean viral load over the simulated individuals at all times with its 95% inter quantile range. Additionally, we provided the distribution of several viral dynamic metrics, namely:

- the area under viral load curve,
- the peak and time to peak viral load
- the duration of the clearance stage, calculated as the time interval between the peak viral load and the time to undetectable viral load
- the duration of the acute phase, calculated as the time between the first and the last detectable viral load (32).

### Parameter estimation

All parameters were estimated by computing the maximum-likelihood estimator using the stochastic approximation expectation-maximization (SAEM) algorithm implemented in Monolix Software 2020R1 (33,34). Standard errors and the likelihood were computed by importance sampling.

## Acknowledgement

We would like to thank everyone in the CEA and at Pasteur Institute that have helped for data collection. We thank Alan Perelson for helpful discussions.

## Supplementary material

**S1 Fig.**
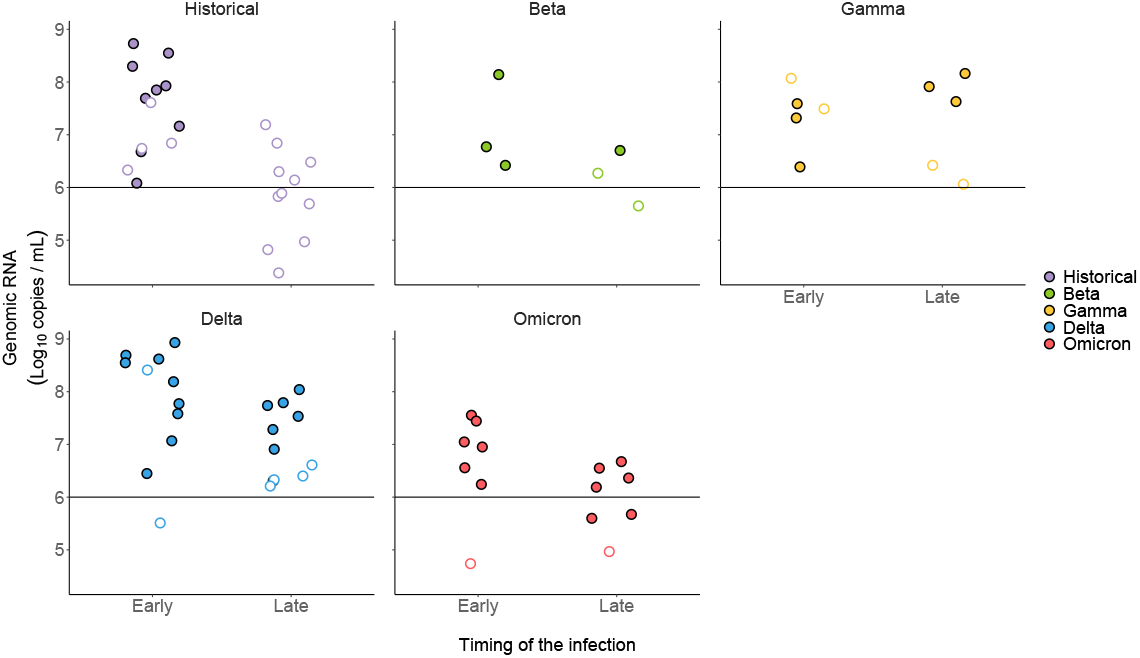
Relationship between genomic RNA and infectious titers. Undetectable infectious titers are depicted as empty circles. The timings early and late correspond to swab sampled at 2, 3 or 4 days post infection and 5 or 7 days post infection respectively.

**S2 Fig.**
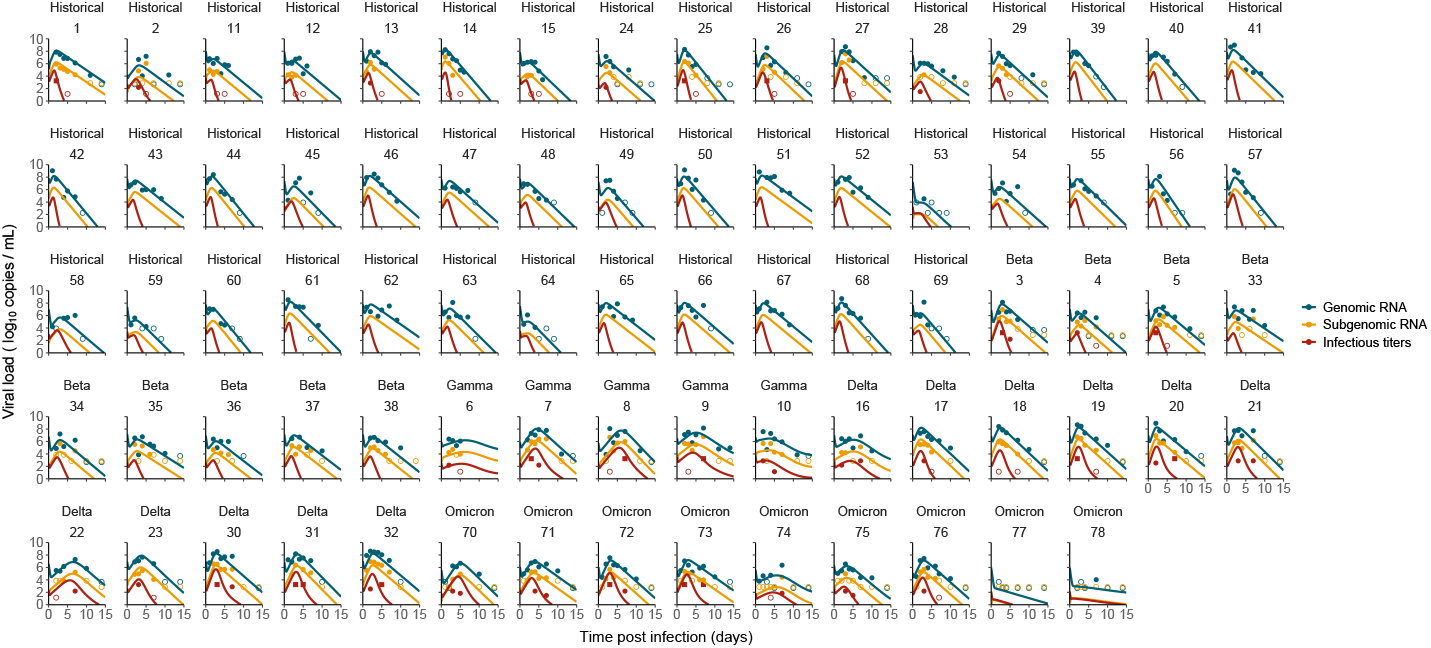
Individual fit of genomic RNA, subgenomic RNA and infectious titers in all animals. Undetectable values are represented as empty dots. Values above the upper limit of quantification are represented as squares.

**S3 Fig.**
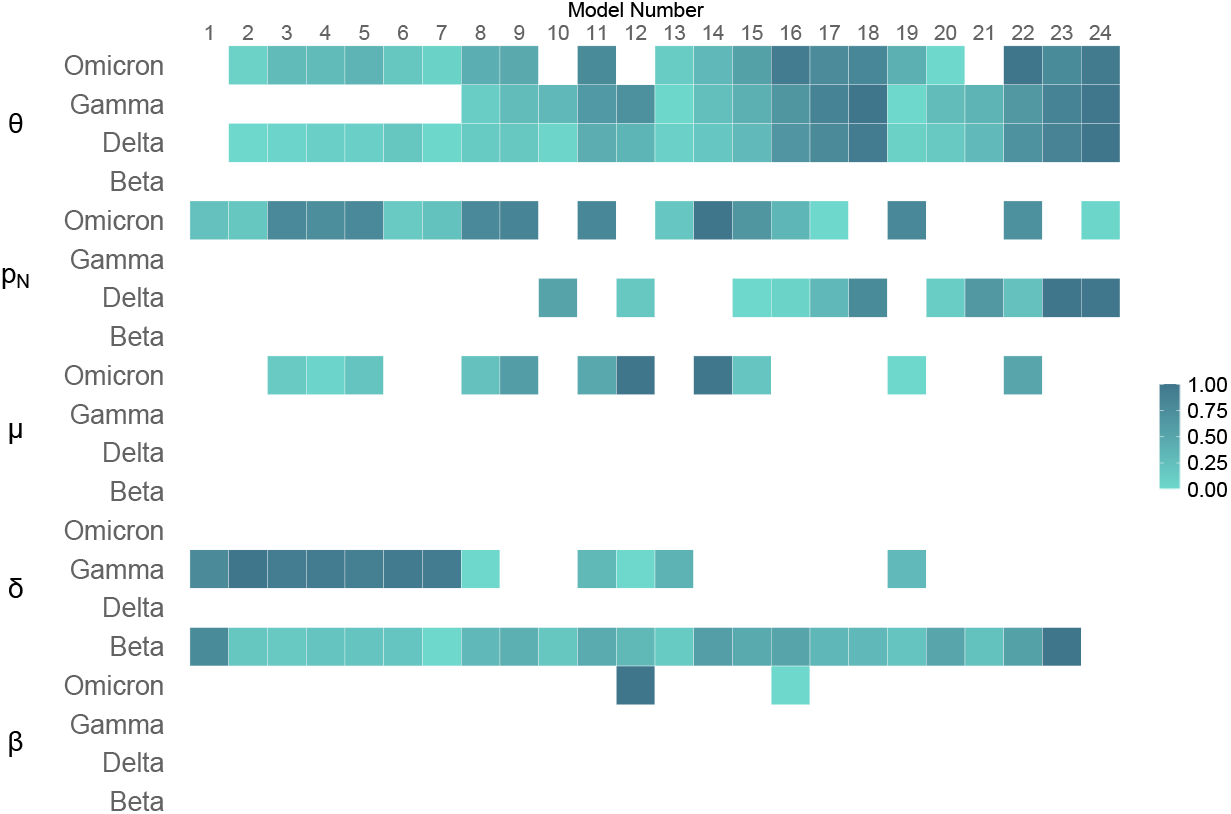
Sensitivity analysis on the covariate selection algorithm. We performed a sensitivity analysis on our best model. The model IDs are represented on top, as described in S3 Table. The scale represents the magnitude of the covariate effect rescaled for each row with 0 being the minimum value and 1 the maximum. Empty tiles indicate that no covariates were selected for this variant-parameter relationship.

**S4 Fig.**
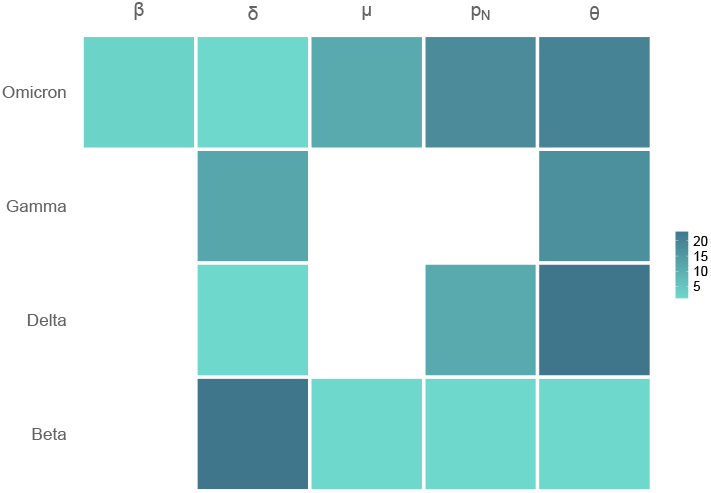
Consistency of the covariate selection algorithm. We represent the number of times a covariate was found on a variant-parameter relationship across all 24 models. Empty tiles indicate that no covariates were found for this variant-parameter relationship.

**S1 Table.** Characteristics of the 78 animals analysed. Descriptive statistics of the animals calculated on the raw data.

| Strains | Number of animals | Mean weight (kg) | Mean peak viral load (log<sub>10</sub> copies/mL) | Mean peak PFU (log<sub>10</sub> PFU/mL) | Mean time to first undetectable viral load | Mean time to first undetectable PFU |
| --- | --- | --- | --- | --- | --- | --- |
| Historical | 44 | 3.7 | 7.6 | 2.3 | 8 | 4 |
| Beta | 9 | 4.9 | 7.1 | 3.2 | 10 | 6 |
| Gamma | 5 | 4.2 | 7.8 | 3 | 14 | 5 |
| Delta | 11 | 3.6 | 8.1 | 2.9 | 12 | 5 |
| Omicron | 9 | 4.6 | 6.4 | 2.4 | 12 | 7 |

**S2 Table.** Estimates of the population parameter and covariate effects for the best model. **The standard error for the *R*_0_ parameters were calculated using the delta method.

| *Population parameters (unit)* | Fixed effect (RSE%) | SD of random effect (RSE%) |
| --- | --- | --- |
| $\beta$ (copie.d<sup>-1</sup>) | $1.85 \times 10^{-5}$ (33) | - |
| $p$ (copies.cell<sup>-1</sup>.d<sup>-1</sup>) | $9.44 \times 10^5$ (40) | 0.61 (17) |
| $\delta$ (d<sup>-1</sup>) | 1.38 (6) | 0.2 (20) |
| $f$ (unitless) | $1.36 \times 10^{-3}$ (19) | - |
| $\mu$ (unitless) | $1.98 \times 10^{-4}$ (47) | - |
| $\theta$ (unitless) | 0.19 (45) | 0.32 (103) |
| *Covariate model* | Covariate effect (RSE%) | p-value |
| Beta on $\delta$ | -0.357 (32) | 0.00201 |
| Gamma on $\theta$ | 8.39 (15) | $< 10^{-6}$ |
| Delta on $p$ | 0.554 (50) | 0.047 |
| Delta on $\theta$ | 5.49 (15) | $< 10^{-6}$ |
| Omicron on $p$ | -2.82 (19) | $< 10^{-6}$ |
| Omicron on $\mu$ | 2.7 (23) | $1.98 \times 10^{-5}$ |
| Omicron on $\theta$ | 5.04 (18) | $< 10^{-6}$ |
| *Basic reproductive number* | Value (RSE%)** |  |
| $R_0$ | 3.1 (19) | |
| $R_{0\beta}$ | 4.5 (20) | |
| $R_{0\gamma}$ | 3.1 (19) | |
| $R_{0\delta}$ | 5.4 (34) | |
| $R_{0\omicron}$ | 2.8 (28) | |
| *Residual errors* | Value (RSE%) |  |
| $\sigma_{\text{Genomic RNA}}$ (log<sub>10</sub> copies/mL) | 0.98 (4) | |
| $\sigma_{\text{Subgenomic RNA}}$ (log<sub>10</sub> copies/mL) | 0.89 (6) | |
| $\sigma_{\text{Infectious titers}}$ (log<sub>10</sub> PFU/mL) | 1.79 (14) | |

**S3 Table.** Sensitivity analysis on the delayed immune response. Using the best structural model (i.e. Model 1 including an effect on the infectious ratio) we tested several delays for the immune response to take place and performed the covariate search algorithm on all models.

| Model ID | Number of transfer compartments $F$ | Delay (days) | Transfer rate parameter $g$ (d<sup>-1</sup>) | BIC before COSSAC | BIC after COSSAC (ΔBIC) |
| --- | --- | --- | --- | --- | --- |
| 1 | 5 | 1 | 5 | 2451 | 2429 (-22) |
| 2 | 5 | 2 | 2.5 | 2409 | 2384 (-25) |
| 3 | 5 | 3 | 1.666666667 | 2405 | 2374 (-31) |
| 4 | 5 | 4 | 1.25 | 2408 | 2373 (-35) |
| 5 | 5 | 5 | 1 | 2408 | 2375 (-33) |
| 6 | 5 | 6 | 0.833333333 | 2409 | 2379 (-30) |
| 7 | 10 | 1 | 10 | 2432 | 2409 (-23) |
| 8 | 10 | 2 | 5 | 2409 | 2373 (-36) |
| 9 | 10 | 3 | 3.333333333 | 2410 | 2361 (-49) |
| 10 | 10 | 4 | 2.5 | 2411 | 2381 (-30) |
| 11 | 10 | 5 | 2 | 2413 | 2366 (-47) |
| 12 | 10 | 6 | 1.666666667 | 2414 | 2367 (-47) |
| 13 | 20 | 1 | 20 | 2426 | 2402 (-24) |
| 14 | 20 | 2 | 10 | 2409 | 2363 (-46) |
| 15 | 20 | 3 | 6.666666667 | 2411 | 2360 (-51) |
| 16 | 20 | 4 | 5 | 2414 | 2377 (-37) |
| 17 | 20 | 5 | 4 | 2416 | 2379 (-37) |
| 18 | 20 | 6 | 3.333333333 | 2417 | 2381 (-36) |
| 19 | 30 | 1 | 30 | 2424 | 2397 (-27) |
| 20 | 30 | 2 | 15 | 2408 | 2377 (-31) |
| 21 | 30 | 3 | 10 | 2413 | 2385 (-28) |
| 22 | 30 | 4 | 7.5 | 2417 | 2363 (-54) |
| 23 | 30 | 5 | 6 | 2419 | 2380 (-39) |
| 24 | 30 | 6 | 5 | 2420 | 2393 (-27) |

